# Distinct value encoding in striatal direct and indirect pathways during adaptive learning

**DOI:** 10.1101/277855

**Authors:** Christopher H. Donahue, Max Liu, Anatol C. Kreitzer

**Affiliations:** The Gladstone Institutes, San Francisco, CA 94158, USA; Neuroscience Graduate Program, University of California, San Francisco, San Francisco, CA 94158, USA; Departments of Physiology and Neurology, University of California, San Francisco, San Francisco, CA 94158, USA

## Abstract

The striatum is thought to play a central role in action selection and reinforcement, and optogenetic experiments suggest differential roles for direct- and indirect-pathway medium spiny neurons (dMSNs and iMSNs). However, the encoding of value-related information in dMSNs and iMSNs during adaptive decision-making is not well understood. We trained mice on a dynamic foraging task where they had to learn the value of different options based on their recent history of choices and outcomes. Single-cell calcium imaging in dorsomedial striatum revealed that dMSNs and iMSNs were oppositely modulated by the updated value of the different options. Additionally, we found that iMSNs were more active as animals slowed between trials, likely reflecting ongoing changes in motivational state. Together, our results demonstrate that co-activation of dMSNs and iMSNs during action initiation does not simply encode action identity, but instead reflects pathway-specific encoding of movement, motivation, and value information necessary for adaptive decision-making.

## INTRODUCTION

The basal ganglia are thought to play a central role in reinforcement learning and decision-making (Doya, 2000; O’Doherty et al., 2004). Medium spiny neurons (MSNs) in the dorsal striatum encode information relevant to computations that are theorized to occur with learning, such as action and chosen values (Samejima et al., 2005; Lau and Glimcher, 2008; Kim et al., 2009; Ito and Doya, 2009, 2015; Cai et al., 2011). Other studies have highlighted a role for dorsal striatum in representing information about reward expectancy (Lauwereyns et al., 2002) and motivational state (Wang et al., 2013), which can potentially drive the invigoration of movement via dopaminergic signaling (Panigrahi et al., 2015; Dudman and Krakauer, 2017). Together, these works suggest that the dorsal striatum is involved in a broad range of value-related computations relevant to adaptive behavior.

Classical models of basal ganglia function proposed opposing roles of striatal direct and indirect pathway medium spiny neurons (dMSNs and iMSNs) in motor control and disease (Albin, 1989; Delong, 1990). Recent theoretical work has suggested that the two pathways also play opposing roles in reinforcement learning, with dMSNs mediating learning from reward and iMSNs underlying punishment (Frank et al., 2004; Collins and Frank, 2014). Experimental support for these models comes largely from optogenetic experiments, in which activation of dMSNs increased movement and drove reinforcement, whereas activation of iMSNs decreased movement and drove punishment (Kravitz et al., 2010, 2012; Yittri and Dudman, 2016). Pathway-specific stimulation can also bias an animal’s choice in opposing directions, suggesting that dMSNs and iMSNs may oppositely encode information about action value (Tai et al., 2012). However, optogenetic experiments can only reveal so much about the encoding of value-related computations by dMSNs and iMSNs.

To gain better insight into the differential roles of these subpopulations, it is necessary to monitor their activity in behaving animals. Surprisingly, recent recording studies have demonstrated that dMSNs and iMSNs behave similarly during simple motor behaviors (Isomura et al., 2013; Cui et al., 2013; Jin et al., 2014; Klaus et al., 2017). However, the behaviors used in these studies were not designed to examine value-based learning. In a recent Pavlovian conditioning study in which cues were associated with different reward probabilities (Shin et al., 2018), expected reward had a differential effect on dMSN and iMSN responses, opening up the possibility that dMSNs and iMSNs may be dynamically modulated during adaptive decision-making.

Here, we recorded large-scale single-cell calcium signals from identified dMSNs and iMSNs as mice performed a dynamic foraging task where they had to learn to choose the better option based on their recent experience. Both dMSNs and iMSNs were co-active as the animals initiated movements, but distinct signals were identified after an outcome was revealed, which were related to the value of different choice options and the motivation of animals to initiate subsequent trials.

## RESULTS

To study the functional role of direct and indirect pathway MSNs in adaptive value-based decision-making, we trained mice (6 D1-Cre and 6 A2a-Cre) to perform a dynamic foraging task where the probability of receiving a reward between two options varied dynamically across trials. To successfully perform the task, animals had to use information from their recent history of choices and outcomes to determine which option was most likely to lead to reward. The task was designed to maximize our ability to dissociate signals related to movement direction and outcomes, with the goal of determining how dMSNs and iMSNs differentially combine this information to guide future choices.

The animals began each trial by entering a central port, and they were then required to make a decision between left and right ports to receive sucrose reward (Fig. 1a). Rewards were independently assigned to each side at different probabilities (60% vs. 15%), and once assigned, remained available until chosen. A similar task design has been used to study matching behavior (Hernstein, 1961; Sugrue et al., 2004; Lau and Glimcher, 2008; Fonesca et al., 2015) and was employed here to encourage the animals to dynamically sample both options. The location of the high reward probability port underwent un-signaled reversals every 80 trials. Animals learned to rapidly adapt their choice behavior in response to changes in reward contingency (Fig. 1b) and combined information about their recent choices and outcomes to make their decisions (Fig. 1c, see methods). As expected, the animals exhibited robust win-stay, lose-switch behavior (WSLS) (Wilcoxon signed rank test: p<10^−3^) with no detectable differences observed across genotypes (**fig. S1**, Wilcoxon rank sum test: p=0.818).

**Figure 1:**
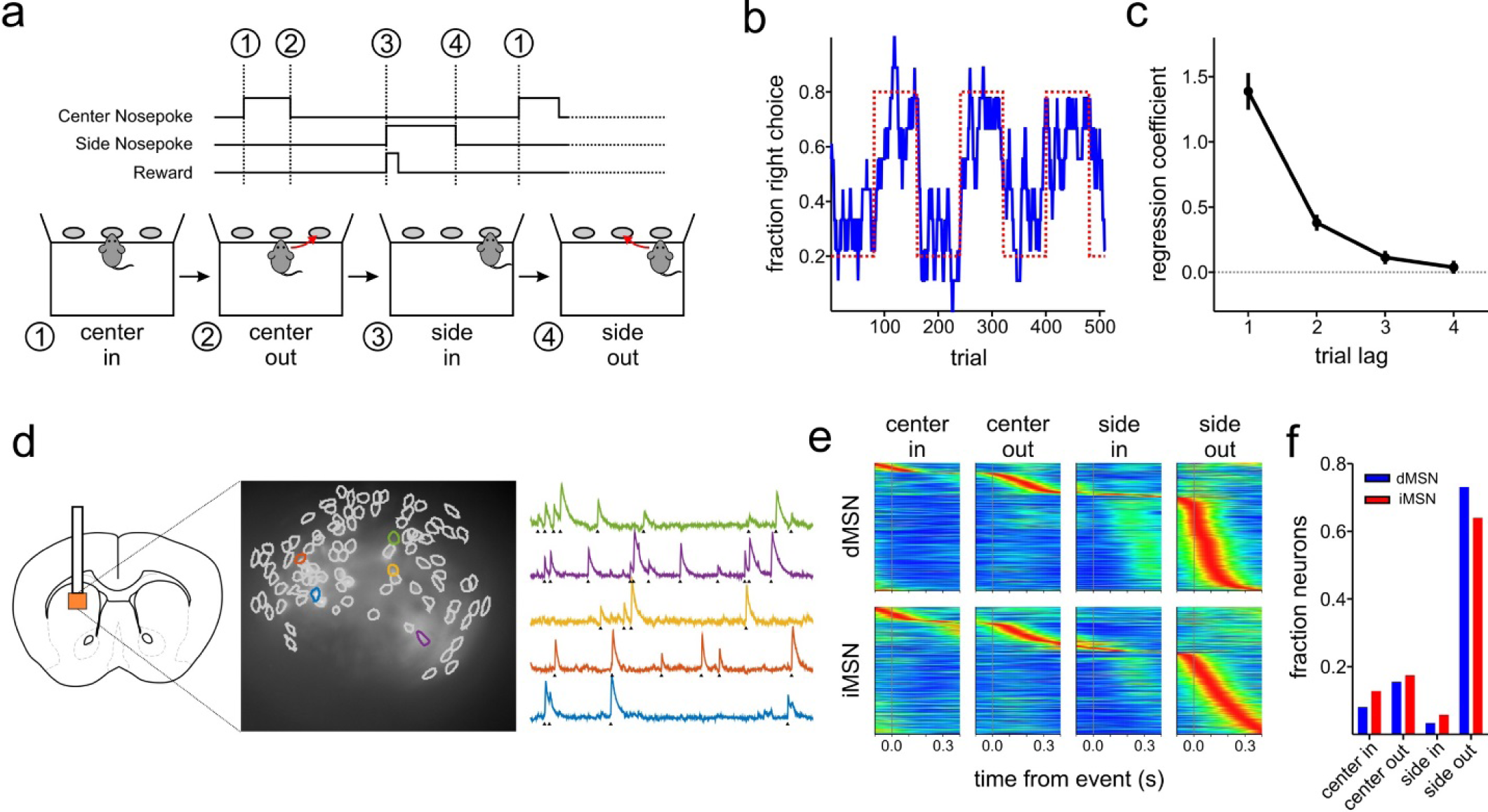
Single cell calcium imaging of dMSNs and iMSNs in a dynamic foraging task. **(a)** Schematic of dynamic foraging task showing sequence of behavioral events. **(b)** Choice behavior from an example animal. The animal adapted its choice behavior (blue, 10-trial moving average) in response to changes in reward contingencies (dashed red line, ratio of right to left side reward probabilities). **(c)** Regression coefficients (averaged across all animals) showing how recent choice and outcome history influences an animal’s upcoming choice. **(d)** (left) gCaMP6m was expressed in the left dorsomedial striatum. A GRIN lens and head-mounted microscope were implanted above the site. (center) Spatial footprints of identified dMSNs obtained from a D1-Cre animal using the CNMF-E algorithm (Zhou et al., 2018). (right) Example calcium traces recorded while the animal performed the dynamic foraging task. Triangles indicate the detected onset time of calcium transients. **(e)** For each neuron, the onset time associated with all calcium transients was aligned to each behavioral event. Neural activity was normalized to each neuron’s maximum response, and neurons were sorted according to the timing of their maximum response. **(f)** The fraction of neurons whose maximum activity is associated with each of the 4 behavioral events shown in (a).

To record single neuron activity from identified dMSNs and iMSNs, we expressed a genetically-encoded calcium indicator (gCaMP6m) in the dorsal striatum of D1-Cre or A2a-Cre mice, and imaged neuronal activity through a GRIN lens with a head-mounted miniature microscope (Fig. 1d and **fig. S2**). We recorded from a total of 360 dMSNs and 448 iMSNs in well-trained animals performing the task. Calcium traces were extracted for each neuron using CNMF-E (Zhou et al., 2018), and event detection was used to align the onset of calcium transients relative to key behavioral events in the task (Fig. 1e and **fig. S3,4a**; see methods). MSN activity was sparse (**fig. S4b**) and largely driven by orienting movements as the animals moved between the ports (Cui et al., 2013; i.e. the center-out or side-out epochs; Fig. 1e). In both the dMSN and iMSN populations, the majority of cells were most active during the side-out movement period (Fig. 1f), a task epoch occurring directly after outcome was revealed, when animals initiated movements back towards the center port to begin a new trial. Given the more robust signaling during the side-out movement, we hypothesized that MSN activity during this task period carried important information beyond simple motor signals. We therefore focused on this epoch to dissociate signals in direct and indirect pathway MSNs related to movement and outcome, and to determine how this information could be combined to eventually influence an animal’s behavior.

We began by characterizing side-out movement-related activity in more detail by deriving a tuning index (see methods) to measure each neuron’s selectivity to either contralateral or ipsilateral movement. The field of view from a D1-Cre animal shown in Fig. 2a depicts dMSNs that were significantly tuned to either contralateral or ipsilateral movements. Neurons that were tuned towards the same direction tended to be clustered closer together (Klaus et al., 2017) than oppositely tuned neurons (see methods, Wilcoxon rank sum test: p=0.003), but no differences were observed between dMSNs and iMSNs (Wilcoxon rank sum test: p=0.610). The two example cells displayed in Fig. 2b demonstrate selectivity for each movement direction. Across both dMSN and iMSN populations, there was an overwhelming bias for movement-selective cells to be preferentially tuned for contralateral movement (Fig. 2c&d, dMSNs: 170/179, 94.5%; iMSNs: 235/253, 92.9%). Interestingly, the overall strength of the contralateral tuning bias across the population was significantly greater in dMSNs compared to iMSNs (Fig. 2e; paired t-test: p=0.005).

**Figure 2:**
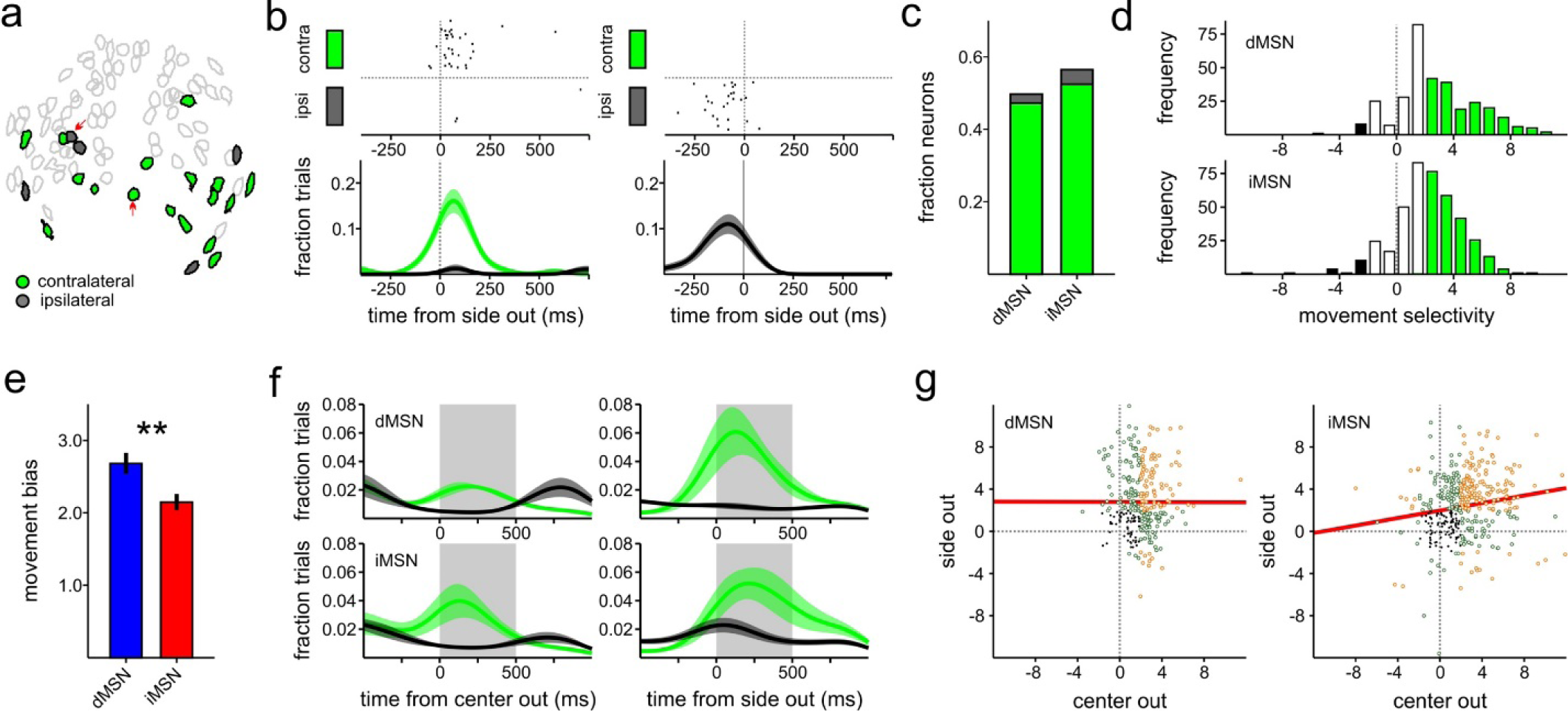
Movement selectivity in dMSNs and iMSNs. **(a)** Spatial footprints of identified dMSNs from an example D1-Cre animal. Filled colors indicate whether an increase in activity was related to contralateral (green) or ipsilateral (black) movement. **(b)** A contralateral-movement preferring cell (left) and ipsilateral-movement preferring cell (right) recorded during the side-out period. (top) Rasters depict the onset time of each calcium transient. (bottom) Density of calcium events for contralateral (green) and ipsilateral (black) movements. Example cells are indicated on the map by arrows in (a). **(c)** Fraction of neurons that were selective for movement during the side-out period (green = contralateral; black = ipsilateral). **(d)** Distribution of movement selectivity indices for dMSNs (top) and iMSNs (bottom). **(e)** Average movement direction bias across all dMSNs and iMSNs. Positive values indicate a preference for contralateral movement. **(f)** Time course of movement-evoked responses aligned to the center-out (left) and side-out (right) movement periods. The density of calcium transients was calculated for each animal and then combined. **(g)** Relationship of movement selectivity indices for each dMSN (left) and iMSN (right) measured during the center-out and side-out periods.

The large differences that we observed when comparing activity during the movement periods (center-out and side-out; Fig. 1e and Fig.1f) led us to question whether individual neurons maintained their directional tuning across both epochs. When we analyzed movement tuning at the population level, we found a significant tuning bias for contralateral movement in both epochs, replicating previous findings (Cui et al., 2013, Fig. 2f). However, when we examined this at the single neuron level, we found that tuning direction was maintained only in the iMSN population (Fig. 2g; r=0.178, p<10^−3^) but not in the dMSN population (r=0.00, p=0.965), suggesting that the directional coding we observed in dMSNs depended upon task context and was not a simple reflection of movement-related information. In further support of this, we found that the movement-selective iMSNs during the center-out period tended to be the same neurons that were movement-selective during the side out period (χ^2^: 31.576, p<10^−7^), while the dMSN population showed no significant overlap (χ^2^=0.499, p=0.480). While recent work has emphasized similar response properties of dMSNs and iMSNs during movement initiation (Cui et al., 2013; Klaus et al., 2017), our results suggest that dMSNs might respond differently when similar movements are made in different contexts.

We next examined how MSNs encoded information about recent outcomes, independent of movement direction (see methods). The iMSNs shown in Fig. 3a were highly selective to either recently rewarded or unrewarded outcomes. Similar to the directionally-selective neurons, outcome-selective neurons were also clustered in space (Wilcoxon rank sum test: p<0.001; see methods), and no differences were detected between dMSNs and iMSNs (Wilcoxon rank sum test: p=0.786). Fig. 3b shows two example neurons that were highly tuned for each outcome. When we looked across all recorded neurons, we found that a significantly larger fraction of iMSNs were outcome-selective (Fig. 3c; iMSNs: 140/448 (31.3%), dMSNs: 68/360 (18.9%); 2-proportion z-test: p<10^−4^). To our surprise, iMSNs demonstrated a strong preference for recently rewarded outcomes (111/140 neurons, 79.3%; binomial test: p<10^−11^), while dMSNs had no bias for either rewarded or unrewarded outcomes (34/68 neurons, 50%; binomial test: p=1.00). We calculated an outcome selectivity index for each cell (Fig. 3d) and found that the mean response of iMSNs was significantly biased towards positive outcomes (Fig. 3e; t-test: p<10^−18^), while dMSNs showed no detectable bias (t-test: p=0.250; paired t-test, iMSNs>dMSNs: p<10^−8^). Although these effects were strongly related to the recent outcome, it is important to note that these signals were best aligned to initiation of movements back to the center port and not to reward delivery (or omission). Thus, they seemed likely to reflect features of the movement that were modulated by the recent outcome.

**Figure 3:**
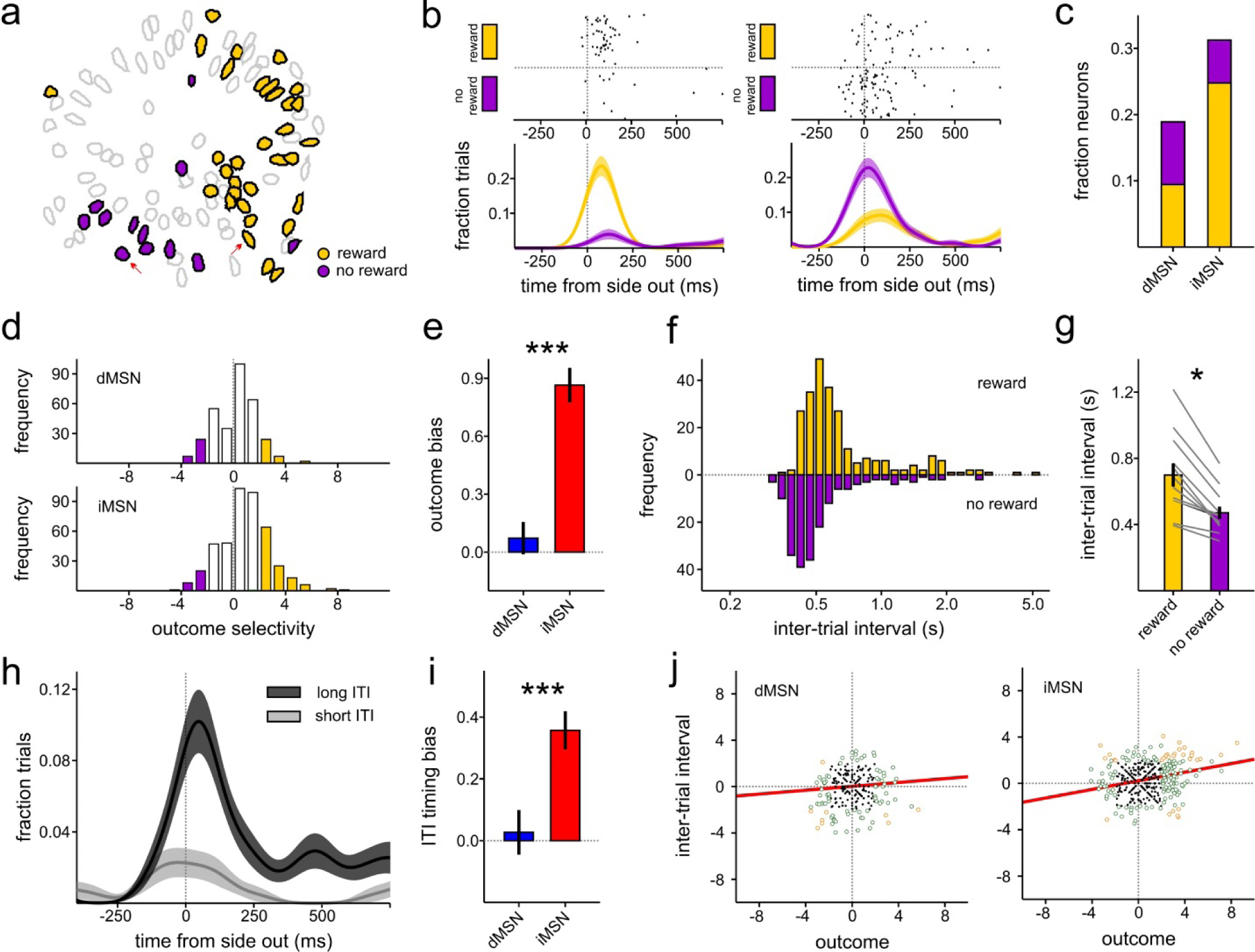
Modulation of dMSNs and iMSNs by outcome and intertrial interval. **(a)** Spatial footprints for identified iMSNs from an example A2a-Cre animal. Filled colors indicate whether an increase in activity was related to rewarded (yellow) or unrewarded (purple) outcomes. **(b)** An example cell that preferred rewarded trials (left) and another that preferred unrewarded trials (right). Cells are indicated on the map by arrows in (a). **(c)** Fraction of neurons that were selective for outcome during the side-out period (yellow = rewarded; purple = unrewarded). **(d)** Distribution of outcome selectivity indices for dMSNs (top) and iMSNs (bottom). **(e)** Average outcome bias in dMSN and iMSN populations. Positive values indicate a preference for rewarded trials. **(f)** Distribution of inter-trial intervals from an example animal with rewarded (yellow) and unrewarded trials (purple) shown separately. **(g)** Average inter-trial interval across all animals for rewarded and unrewarded trials. **(h)** Example iMSN that showed increased activity during movements that preceded longer inter-trial intervals. **(i)** Average bias for ITI timing selectivity across dMSN and iMSN populations. Positive values indicate a preference for longer ITIs. **(j)** Relationship of outcome tuning and ITI tuning for the population of dMSNs (left) and iMSNs (right) measured during the side-out period.

To investigate this further, we examined timing behavior to determine how the inter-trial interval (ITI) was influenced by the recent outcome. Since this was a self-paced task, the time to reengage in the next trial could potentially reflect information about the animal’s current motivational state. The example animal’s behavior in Fig. 3f shows that the ITI was significantly faster following unrewarded trials (paired t-test: p<10^−5^), and this pattern was observed across all animals (Fig. 3g, Wilcoxon rank sum test: p=0.017). This phenomenon, known as the post-reinforcement pause (Ferster and Skinner, 1957), is commonly observed in animals performing self-paced operant tasks and is thought to be related to motivational factors such as the expected size or distance from future rewards.

We next sought to determine whether the bias in positive outcome coding that we observed in the iMSN population was directly related to the post-reinforcement pause. The example iMSN neuron shown in Fig. 3h was more active during side out movements that were associated with slower ITIs. We calculated an ITI selectivity index for each neuron (**fig. S5**; see methods) and found that the iMSN population was significantly more active for slower ITIs (t-test: p<10^−7^) while the dMSN population showed no significant bias for either faster or slower ITIs (Fig. 3i; t-test: p=0.724; paired t-test: iMSNs>dMSNs, p<10^−3^). We next examined the relationship between ITI and outcome tuning and found that outcome and ITI tuning were positively correlated only in iMSNs (Fig. 3j; r=0.186, p<10^−4^) and not in dMSNs (r=0.084, p=0.134), suggesting that iMSNs may play a specific role in the slowing that is associated with the post-reinforcement pause. In further support of this hypothesis, we found that iMSNs showing significant outcome modulation tended to also modulate with ITI (χ^2^=11.29, p<10^−3^), whereas this was not observed in dMSNs (c^2^=1.88, p=0.170).

The dynamic foraging task requires animals to integrate their recent choice and outcome history in order to learn which port is more likely to be associated with reward (Fig. 1c). Since we found that MSNs were clearly modulated by both movement direction and outcome, we next determined whether MSNs combined this information in a way that could guide their future behavior. To investigate this, we compared neural activity on trials where the animals obtained evidence that the contralateral side was better (i.e. unrewarded ipsilateral choices or rewarded contralateral choices) to trials where the animals received evidence that the ipsilateral side was better (Fig. 4a; i.e. rewarded ipsilateral choices or unrewarded contralateral choices). In the example D1-Cre animal shown in Fig. 4b, we found neurons in the same field of view that were significantly modulated by both of these scenarios. In contrast to the movement and outcome-selective neurons, we were unable to detect a tendency for this population to be clustered in space (Wilcoxon rank sum test: p=0.141). The example dMSN shown in Fig. 4c was more active in situations where the animal received feedback that the contralateral side was better. Although there was no difference in the fraction of neurons that were significantly modulated by the combination of choice and outcome in dMSNs and iMSNs (dMSNs: 73/360, 20.3%; iMSNs: 92/448, 20.5%), we found that a significant proportion of dMSNs were biased towards representing evidence that the contralateral side was better (47/73, 64.4%; binomial test: p=0.019) while the iMSN population showed a weak bias in the opposite direction (Fig. 4d; 37/92, 40.2%; binomial test: p=0.076). In support of this, when we derived a tuning index to measure how choice and outcome information was integrated to represent the high value location (Fig. 4e), we found that the mean dMSN and iMSN population responses were biased in opposite directions (Fig. 4f; paired t-test: p<10^−5^). Next, we examined the relationship between choice and outcome tuning at the single neuron level. The outcome tuning of dMSNs was highly predictive of their movement tuning (Fig. 4g&h; r=0.258, p<10^−5^). For example, neurons that were selective for unrewarded (rewarded) trials tended to also be selective for movements that originated from the ipsilateral (contralateral) port. In contrast, these systematic tuning relationships were not present in the iMSN population on a neuron-by-neuron basis (r=0.051, p=0.285). These results indicate that although outcome preference was mixed in dMSNs (Fig. 3e), individual neurons may be commonly tuned along an axis to indicate that recent evidence led to an increase in the value of the contralateral side.

**Figure 4:**
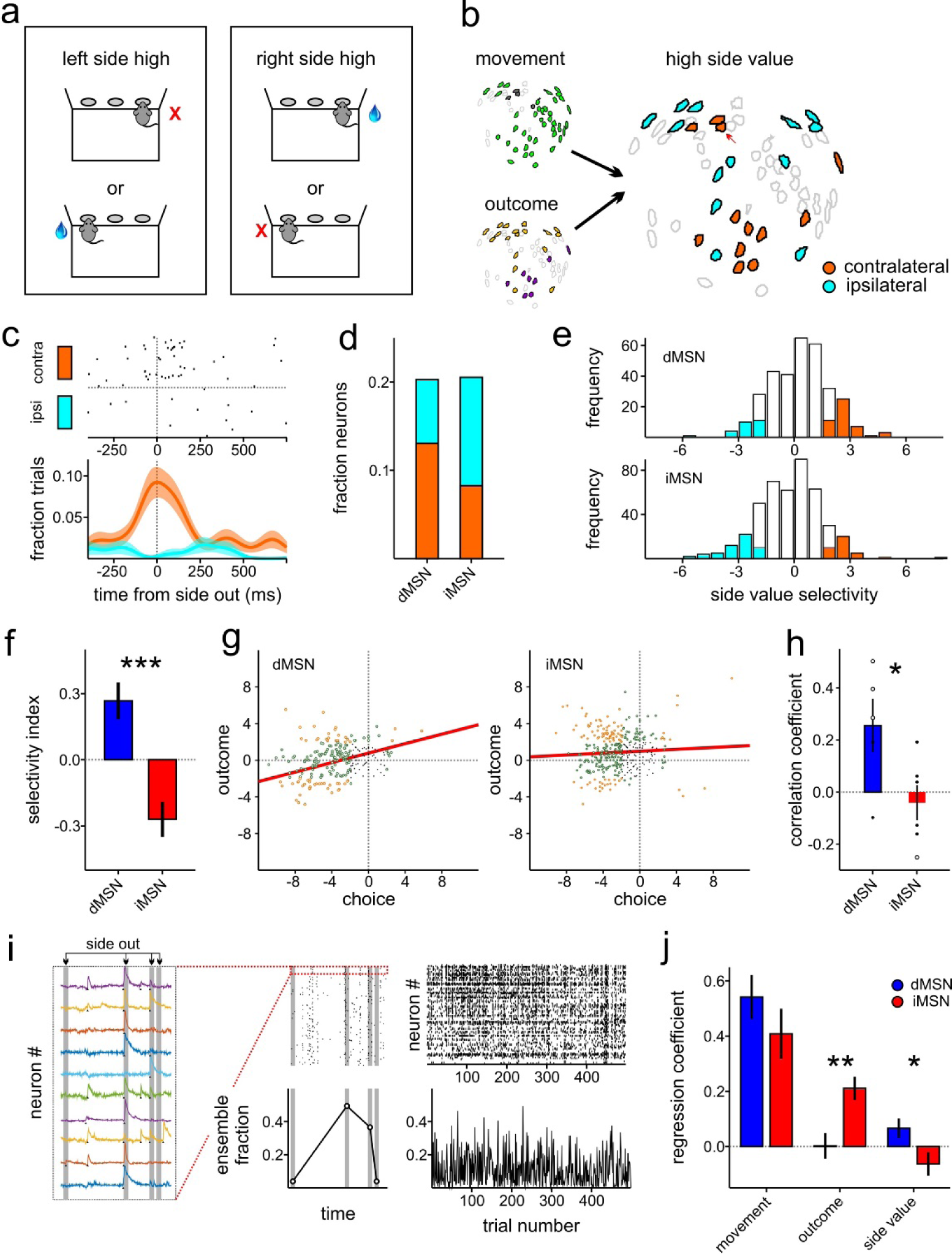
Encoding of side value in dMSNs and iMSNs. **(a)** Information about movement direction and outcome can be combined to indicate the value of a side. **(b)** Spatial footprints for identified dMSNs from an example D1-Cre animal indicating movement, outcome, and side value selectivity. **(c)** Example cell that prefers trials where evidence was in favor of the contralateral side being higher in value. **(d)** Fraction of neurons that were selective for the high-value side during the side-out period (orange = contralateral; cyan = ipsilateral). **(e)** Distribution of high-value location selectivity indices for dMSNs (top) and iMSNs (bottom). **(f)** Average high-value location bias in dMSN and iMSN populations. Positive values indicate a preference for the contralateral side. **(g)** Relationship of outcome tuning and choice tuning for each dMSN (left) and iMSN (right) measured during the side-out period. **(h)** Correlation coefficients associated with neurons shown in (g) plotted for each animal separately. **(i)** Example ensemble analysis. (left) Sample of ten neurons acquired as an animal performed four trials. Gray windows indicate 500ms side-out periods. (center, top) Rasters depict the onset time of calcium transients for all recorded neurons for the same four trials. The highlighted portion corresponds to the example cells in the left panel. (center, bottom) The fraction of the ensemble that is active in each side-out analysis window. (right, top) Raster depicting side-out activity for all neurons and trials recorded from one session. (right, bottom) Fraction of ensemble active during side-out period for each trial. **(j)** Regression coefficients were calculated from the ensemble data from each animal and then combined. The model included terms for movement, outcome, and the location of the high value side.

Finally, we sought to further confirm these findings by leveraging the unique advantage of single neuron calcium imaging to measure the ensemble response of dMSN and iMSN populations on a trial-by-trial basis (Fig. 4i). We analyzed how modulation in the fraction of co-active units varied with either choice (i.e. movement), outcome, or the interaction between choice and outcome (Fig. 4j). When we ran a multiple linear regression model for each animal separately (see methods), we confirmed that the iMSN ensemble response was significantly biased for positive outcomes (Wilcoxon rank sum test: iMSNs>dMSNs, p=0.015). Additionally, dMSN and iMSN populations were oppositely tuned such that more dMSNs were co-active when the animals received evidence that the contralateral port increased in value, and more iMSNs were co-active when evidence favored the ipsilateral port (Wilcoxon rank sum test: p=0.041). Importantly, these ensemble-level analyses were performed on the population response of each animal separately, demonstrating that the effects we previously described at the single-neuron level were a common feature observed across animals. These results are consistent with the possibility that dMSNs and iMSNs might encode action values in an opposing manner as has been suggested by recent optogenetic experiments (Tai et al., 2012). While the single neuron responses in our dataset were too sparse to fit to trial-by-trial estimates of action values, we were able to use the ensemble responses to confirm that our results are consistent with the possibility that direct and indirect pathway MSNs represent opposing action-value representations (**fig. S6**).

## DISCUSSION

Our results demonstrate several clear functional dissociations between dorsal striatal direct and indirect pathway MSNs during value-based decision-making. Recent work using fiber photometry (Cui et al., 2013) and single cell calcium imaging during freely moving behavior (Klaus et al., 2017) has emphasized coordinated activity amongst dMSNs and iMSNs, demonstrating that these two subpopulations similarly increase activity when animals initiate contralateral movements. When we analyzed movement-evoked activity during a decision-making task, we also observed concurrent activity in dMSNs and iMSNs that was strongly biased for contralateral movement. However, when we decomposed these signals into factors related to outcome and movement direction, we were able to reveal opposing features of dMSN and iMSN activity. This was made possible by combining single-cell recording with a behavioral task designed to elicit highly stochastic behavior, giving us the statistical power to dissociate movement direction and outcome over many trials. Prior efforts to discern functional differences in dMSN and iMSN activity may have been hindered by limitations inherent to calcium imaging, given the sparse responses we observed in MSNs. Indeed, recent studies using more sensitive measures of neural activity such as whole cell recording and electrophysiology have revealed distinct differences between these populations in sensory processing (Sippy et al., 2015) and in reward expectation (Shin et al., 2018).

The new contributions of our study rest on two main findings. First, we showed that dMSNs and iMSNs were oppositely modulated by changes in side value, with dMSN activity increasing when recent evidence favored an increase in the relative value of the contralateral side, and iMSN activity increasing when it favored an increase in the relative value of the ipsilateral side. Second, we found that iMSNs may play a specific role in slowing during the inter-trial interval, which was especially pronounced following rewarded outcomes.

### Action value in the dorsal striatum

The striatum has been theorized to play a central role in computations involved in reinforcement learning (Sutton and Barto, 1998), and electrophysiological recordings from unidentified MSNs suggest that the dorsal striatum encodes trial-by-trial representations of action value (Samejima et al., 2005; Lau and Glimcher, 2008; Kim et al., 2009). Optogenetic stimulation of dMSNs and iMSNs imply that they may oppositely encode action values (Tai et al., 2012), but recordings from identified dMSN and iMSN striatal populations during decision-making tasks have not yet been reported. We found that dMSNs were more active after the animals received evidence that the relative value of the contralateral port increased, and iMSNs were more active following evidence that the relative value of ipsilateral port increased. Indeed, when we modeled the behavior with a simple reinforcement learning model, we found that ensemble responses in dMSNs and iMSNs were oppositely modulated by relative differences in action values.

Interestingly, while we found that dMSN and iMSN populations were oppositely modulated, choice and outcome tuning at the single-neuron level was only correlated in dMSNs (Fig 4g). For example, dMSN neurons that were more active after the animal exited the ipsilateral (or contralateral) port tended to also signal that the trial was unrewarded (or rewarded). This relationship was random in iMSNs. This raises the intriguing possibility that there may be a highly specific pattern of choice and outcome inputs to dMSNs in particular, which may point to a unique role in transmitting this information in downstream basal ganglia circuitry. This will be an interesting avenue for future exploration.

The striatum is a highly lateralized structure, displaying a strong preference for motor responses (Cui et al., 2013; Klaus et al., 2017) or expectation of reward (Lauwereyns et al., 2002) arising from the execution of movements made in a contralateral direction with respect to the recording site. Our dynamic foraging task was designed such that the value associated with making either a contralateral or ipsilateral movement to one of the ports was dynamically modulated across trials. We employed this design to take advantage of the natural spatial bias in the striatum, assuming that if MSNs fundamentally care about spatial information, we would be more likely to detect any value-based modulation across space. However, the extreme laterality of the signals we recorded was also a disadvantage since so few neurons were active whenever the animals moved in the ipsilateral direction. Thus, it would be extremely valuable for future studies to employ alternative task designs where actions are not lateralized, such as requiring animals to make decisions by manipulating a lever.

### Role of iMSNs in slowing

Our finding that iMSNs were more active following rewarded trials was unexpected and contrary to predictions of optogenetic (Kravitz et al., 2012) and theoretical (Frank et al., 2004; Collins and Frank, 2014) work. In an electrophysiological study from optogenetically identified dMSNs and iMSNs in a Pavlovian conditioning task, dMSNs tended to increase activity when the animal expected higher reward while iMSNs displayed the opposite effect (Shin et al., 2018). We can explain this apparent discrepancy by noting that the signals we report occurred well after outcome delivery and consumption of reward, and were best aligned to the initiation of movement back towards the central port. Instead of being related to reward processing, we suggest that these iMSN responses were related to the motor slowing that occurred following rewarded trials. In support of this hypothesis, we found that iMSNs were significantly more likely to respond to movements associated with long versus short ITI trials, and that the population of outcome-selective iMSNs tended to be the same population that was selective to ITI timing. Indeed, optogenetic stimulation of striatal iMSNs slows or suppresses movement (Kravitz et al., 2010). Our results suggest that the indirect pathway might play a role in slowing during natural behaviors, potentially reflecting ongoing changes in an animal’s motivational state.

We observed longer inter-trial intervals following reward, consistent with what is known as the ‘post-reinforcement pause’. This phenomenon has been described across a wide range of species in self-paced operant tasks (Ferster and Skinner, 1957; Schlinger et al., 2008). It is most pronounced in fixed ratio tasks where the animals are required to make a fixed number of operant responses to receive reward, and it tends to systematically increase with higher ratio requirements. It is thought that the pause reflects a momentary decrease in motivation since reward delivery indicates that the next reward is now more distant or will require more effort to achieve. Interestingly, our task used a design where ports were baited with reward such that the overall reward probability progressively increased following each unrewarded trial and was actually lowest following reward. This may have played a role in the robust slowing that we observed after reward, and could explain apparent differences between ours and other studies (Wang et al., 2013). Other potential differences might be attributed to the fact that our task was entirely self-paced (animals were allowed to reengage the next trial whenever they wanted), while other behaviors often impose a fixed ITI and measure reengagement as the time to respond to a trial initiation cue (Wang et al., 2013). One interesting possibility is that dopaminergic projections to the striatum, which reflect moment-by-moment changes in state value (Hamid et al., 2016), potentially influence the post-reinforcement pause. iMSNs may inversely reflect motivational value via dopaminergic modulation at D2 receptors, serving as a potential way for dopamine to interface with the motor system and produce the behavioral effects related to momentary shifts in motivation.

## METHODS

### Subjects

Male and female transgenic mice (aged 3-6 months) expressed Cre recombinase under control of the dopamine D1 receptor (D1-Cre) to image direct pathway MSNs or adenosine A2a receptor (A2a-Cre) to image indirect pathway MSNs. Mice were maintained on a 12h/12h light/dark cycle and fed ad libitum. Mice were water deprived for 24 hours prior to behavioral training and weight was maintained at >85% of pre-deprivation weight. All procedures were approved by the University of California San Francisco Institutional Animal Care and Use Committee.

### Surgical procedures

Mice were anaesthetized with isoflurane and place in a stereotaxic frame (Kopf). All animals underwent two surgical procedures. In the first procedure, 1uL of GCaMP6m (AAV1-Syn-Flex-GCaMP6m; titer: 3.78 × 10^12^/ml; University of Pennsylvania Vector Core) was injected into the dorsomedial striatum (AP: +0.8, ML: −1.5, DV: −2.5) of D1-cre or A2a-cre mice to achieve selective expression in direct or indirect pathway MSNs. A gradient index lens (GRIN; Inscopix; either 0.5 mm or 1.0 mm) was implanted 200µm above the injection site (DV: −2.3). After a two-week recovery period, mice underwent a second procedure to affix a baseplate above the lens for mounting a miniature fluorescence microscope (Inscopix). Buprenorphine HCl (0.1 mg/kg, intraperitoneal injection) and Ketoprofen (5 mg/kg, subcutaneous injection) were used for postoperative analgesia. Mice were allowed to recover for 1 week before training began.

### Behavior

#### Apparatus

All experiments took place in acrylic custom-built operant chambers (4” × 7”). Each chamber was fitted with three custom-built nose pokes containing built-in infrared emitters and receivers (Sparkfun) and a white LED (Sparkfun). Sucrose was delivered through a stainless steel tube located inside each nose poke and reward delivery was controlled with a solenoid valve (NResearch). The timing of all task events was controlled by a microcontroller (Mbed) and custom software (Statescript, Spike Gadgets).

#### Behavioral Task

In the dynamic foraging task, mice were required to initiate a trial by poking their nose into a central port and hold for a period of 150-300ms (exponentially distributed). At the end of the hold period, an LED illuminated inside the nose poke signaling that a decision could be made. Mice were then free to enter either the left or the right nose poke to collect reward (2µL/reward) via tubing located inside of the chosen port. Reward was delivered immediately upon entering the chosen port.

At the beginning of each trial, rewards were independently assigned to each side according to different probabilities (15% for the low side and 60% for the high side). Once assigned to a port, a reward remained available until the animal eventually chose the respective port (Sugrue et al., 2004; Lau and Glimcher, 2008; Fonseca et al., 2015). After 80 trials, the contingencies were reversed. The animals completed an average of 480.0 trials per recording session. The task was self-paced and we did not impose a fixed inter-trial interval.

### Data Analysis

#### Behavioral modeling

To determine how an animal’s choice was affected by its recent choice and reward history, we employed the following logistic regression model:

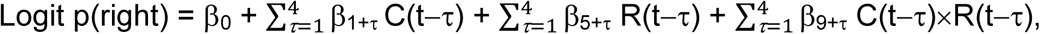

where C(t) is the choice on trial t (1 if right, −1 if left), and R(t) is the outcome on trial t (1 if rewarded, −1 if unrewarded). Coefficients corresponding to the interaction of choice and reward history, C(t-τ)×R(t-τ), are plotted in Fig. 1c.

We fit the following reinforcement learning model to derive trial-by-trial estimates of action-values:

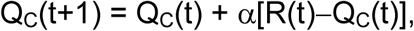

where Q_C_(t) is the value of the chosen port on trial t, R(t) indicates whether a reward was received on trial t (1 = rewarded, 0 = unrewarded), and α is the learning rate. Action selection was modeled with the following equation:

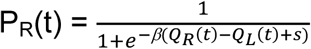

where P_R_ is the probability that the animal chooses the rightward port on trial t, β is the inverse temperature, and *s* is a term accounting for side biases. The three model parameters (a, b, and *s*) were fit for each session separately using a maximum-likelihood procedure.

#### Processing of calcium signals and detection of calcium transients

Images were acquired at 15 frames per second using nVista HD (Inscopix). Recording sessions lasted about one hour. After acquisition, we spatially down-sampled each video by a factor of 4 and corrected for motion-induced artifacts (Mosaic, Inscopix). Cell segmentation was performed with the CNMF-E method (Zhou et al., 2018).

To extract the onset times of calcium events for each neuron, we first found the video frame associated with the peak of each transient (findPeaks.m function in Matlab). To find the onset time associated with each event, we located the frame prior to each peak where the slope of the calcium signal first exceeded zero. Only events whose magnitude was greater than the mean + 1 standard deviation (measured as the difference between the amplitudes at the peak and onset time) were saved.

We aligned all neural data based on four key time points in each trial. The center-in time was defined as the moment the animals entered the central port to start a new trial. The center-out time marked the moment the animal exited the central port. The side-in time was defined as the entry time to the chosen port, and the side-out time was defined as the time that the animals exited the chosen port. The inter-trial interval was defined as the time between side-out and the next trial’s center-in.

#### Calculation of selectivity indices

To calculate single-cell selectivity indices for choice and outcome, we balanced each dataset so that there were an equal number of trials in each of the four movement X outcome categories (i.e. contralateral-rewarded, contralateral-unrewarded, ipsilateral-rewarded, and ipsilateral-unrewarded). For each trial, we determined whether or not a calcium event was detected in a 0-500ms window relative to each behavioral event. The movement selectivity index was calculated by comparing the fraction of contralateral vs. ipsilateral trials (2-proportion z-test) in which a calcium event was detected. The balancing procedure assured that there were an equal number of rewarded and unrewarded outcomes in each group. The outcome selectivity index was calculated using the same subset of trials, but this time comparing rewarded and unrewarded outcomes. To calculate selectivity index corresponding to side value, we used the same balancing procedure, but this time grouped contralateral-rewarded and ipsilateral-unrewarded trials and compared to ipsilateral-unrewarded and contralateral-rewarded trials, thereby balancing by both movement direction and outcome across groups. To calculate the selectivity index related to the inter-trial interval (ITI), we split all trials according to whether the ITI was longer or shorter than the median ITI. Next, we balanced conditions by outcome type and ITI and computed the index using the same procedures described above.

#### Clustering analysis

To determine whether neurons tuned to the same category were clustered in space, we first calculated the mean pair-wise distance between each neuron that was tuned to the same category. For example, we found the mean distance between each neuron that showed significant tuning for contralateral movement, and then repeated this separately for all neurons that were tuned for ipsilateral movement. We combined these distances to determine a mean *within-category* distance. Next, we found the mean *between-category* distance by calculating the distance between each pair that was tuned for opposing movements (i.e. contralateral vs. ipsilateral). We then calculated a ratio of these two values in order to determine whether neurons were more clustered than expected by chance. The ratio of randomly located cells should be equal to 1, and clustered cells should lead to ratios less than 1. To determine statistical significance, we assigned each neuron to a random category and derived a null distribution by calculating the distance ratio on the shuffled datasets 10000 times. The actual ratio was then compared to the null distribution.

#### Ensemble analysis

For each trial, we found the fraction of neurons that were co-active within the 500ms analysis window following the onset of the side-out movement. We used the following multiple linear regression model:

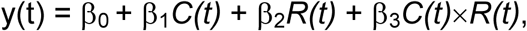

where y(t) is the fraction of neurons that was co-active on trial t, *C(t)* is the chosen port (1 = contralateral, −1 = ipsilateral), *R(t)* is the recently received outcome (1 = reward, −1 = no reward), and *C(t)* × *R(t)* is the interaction term determining the high value location. We calculated standardized regression coefficients from the model for each animal separately.

To determine whether ensemble responses were correlated with relative differences in action value estimates, we used the following multiple linear regression model on the same 500ms analysis window following side-out movement:

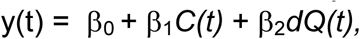

where C(t) is the chosen port on trial t (1 = contralateral, −1 = ipsilateral), and dQ(t) is the difference in action value estimates (Q_R_(t)-Q_L_(t)) derived from a simple reinforcement learning model.

## CONTRIBUTIONS

C.H.D. and A.C.K. designed the experiments. C.H.D. and M.L. performed the experiments. C.H.D. analyzed the data. All authors wrote the manuscript.

## ACKNOWLEDGEMENTS

We thank D. Schulte and B. Margolin for technical assistance, D. Lee for comments on this manuscript, and the Kreitzer and Berke labs for helpful discussion. This research was supported by the NIH (F32 MH107206 to C.H.D. and U01 NS094342 to A.C.K.).

## REFERENCES

Albin, R. L., Young, A. B. & Penney, J. B. The functional anatomy of basal ganglia disorders. Trends Neurosci 12, 366–375 (1989).

Cai, X., Kim, S. & Lee, D. Heterogeneous coding of temporally discounted values in the dorsal and ventral striatum during intertemporal choice. Neuron 69, 170–182, doi:10.1016/j.neuron.2010.11.041 (2011).

Collins, A. G. & Frank, M. J. Opponent actor learning (OpAL): modeling interactive effects of striatal dopamine on reinforcement learning and choice incentive. Psychol Rev 121, 337–366, doi:10.1037/a0037015 (2014).

Cui, G. et al. Concurrent activation of striatal direct and indirect pathways during action initiation. Nature 494, 238–242, doi:10.1038/nature11846 (2013).

DeLong, M. R. Primate models of movement disorders of basal ganglia origin. Trends Neurosci 13, 281–285 (1990).

Doya, K. Complementary roles of basal ganglia and cerebellum in learning and motor control. Curr Opin Neurobiol 10, 732–739 (2000).

Dudman, J. T. & Krakauer, J. W. The basal ganglia: from motor commands to the control of vigor. Curr Opin Neurobiol 37, 158–166, doi:10.1016/j.conb.2016.02.005 (2016).

Ferster, C. B. Schedules of reinforcement. (Appleton-Century-Crofts, 1957).

Fonseca, M. S., Murakami, M. & Mainen, Z. F. Activation of dorsal raphe serotonergic neurons promotes waiting but is not reinforcing. Curr Biol 25, 306–315, doi:10.1016/j.cub.2014.12.002 (2015).

Frank, M. J., Seeberger, L. C. & O’Reilly RC., By carrot or by stick: cognitive reinforcement learning in parkinsonism. Science 306, 1940–1943, doi:10.1126/science.1102941 (2004).

Hamid, A. A. et al. Mesolimbic dopamine signals the value of work. Nat Neurosci 19, 117–126, doi:10.1038/nn.4173 (2016).

Herrnstein, R. J. Relative and absolute strength of response as a function of frequency of reinforcement. J. Exp. Anal. Behavior 4,267–272 (1961).

Isomura, Y. et al. Reward-modulated motor information in identified striatum neurons. J Neurosci 33,10209–10220, doi:10.1523/JNEUROSCI.0381-13.2013 (2013).

Ito, M. & Doya, K. Validation of decision-making models and analysis of decision variables in the rat basal ganglia. J Neurosci 29, 9861–9874, doi:10.1523/JNEUROSCI.6157-08.2009 (2009).

Ito, M. & Doya, K. Distinct neural representation in the dorsolateral, dorsomedial, and ventral parts of the striatum during fixed- and free-choice tasks. J Neurosci 35, 3499–3514, doi:10.1523/JNEUROSCI.1962-14.2015 (2015).

Jin, X., Tecuapetla, F. & Costa, R. M. Basal ganglia subcircuits distinctively encode the parsing and concatenation of action sequences. Nat Neurosci 17, 423–430, doi:10.1038/nn.3632 (2014).

Kim, H., Sul, J. H., Huh, N., Lee, D. & Jung, M. W. Role of striatum in updating values of chosen actions. J Neurosci 29, 14701–14712, doi:10.1523/JNEUROSCI.2728-09.2009 (2009).

Klaus, A et al. The Spatiotemporal Organization of the Striatum Encodes Action Space. Neuron 95, 1171–1180 e1177, doi:10.1016/j.neuron.2017.08.015 (2017).

Kravitz, A. V. et al. Regulation of parkinsonian motor behaviours by optogenetic control of basal ganglia circuitry. Nature 466, 622–626, doi:10.1038/nature09159 (2010).

Kravitz, A. V., Tye, L. D. & Kreitzer, A. C. Distinct roles for direct and indirect pathway striatal neurons in reinforcement. Nat Neurosci 15, 816–818, doi:10.1038/nn.3100 (2012).

Lau, B. & Glimcher, P. W. Value representations in the primate striatum during matching behavior. Neuron 58, 451–463, doi:10.1016/j.neuron.2008.02.021 (2008).

Lauwereyns, J., Watanabe, K., Coe, B. & Hikosaka, O. A neural correlate of response bias in monkey caudate nucleus. Nature 418, 413–417, doi:10.1038/nature00892 (2002).

O’Doherty, J. et al. Dissociable roles of ventral and dorsal striatum in instrumental conditioning. Science 304, 452–454, doi:10.1126/science.1094285 (2004).

Panigrahi, B. et al. Dopamine Is Required for the Neural Representation and Control of Movement Vigor. Cell 162, 1418–1430, doi:10.1016/j.cell.2015.08.014 (2015).

Samejima, K., Ueda, Y., Doya, K. & Kimura, M. Representation of action-specific reward values in the striatum. Science 310, 1337–1340, doi:10.1126/science.1115270 (2005).

Schlinger, H. D., Derenne, A. & Baron, A. What 50 years of research tell us about pausing under ratio schedules of reinforcement. Behav Anal 31, 39–60 (2008).

Shin, J. H., Kim, D. & Jung, M. W. Differential coding of reward and movement information in the dorsomedial striatal direct and indirect pathways. Nat Commun 9, 404, doi:10.1038/s41467-017-02817-1 (2018).

Sippy, T., Lapray, D., Crochet, S. & Petersen, C. C. Cell-Type-Specific Sensorimotor Processing in Striatal Projection Neurons during Goal-Directed Behavior. Neuron 88, 298–305, doi:10.1016/j.neuron.2015.08.039 (2015).

Sugrue, L. P., Corrado, G. S. & Newsome, W. T. Matching behavior and the representation of value in the parietal cortex. Science 304, 1782–1787, doi:10.1126/science.1094765 (2004).

Sutton, R. S. & Barto, A. G. Reinforcement learning: an introduction. (MIT Press, 1998).

Tai, L. H., Lee, A. M., Benavidez, N., Bonci, A. & Wilbrecht, L. Transient stimulation of distinct subpopulations of striatal neurons mimics changes in action value. Nat Neurosci 15, 1281–1289, doi:10.1038/nn.3188 (2012).

Wang, A. Y., Miura, K. & Uchida, N. The dorsomedial striatum encodes net expected return, critical for energizing performance vigor. Nat Neurosci 16, 639–647, doi:10.1038/nn.3377 (2013).

Yttri, E. A. & Dudman, J. T. Opponent and bidirectional control of movement velocity in the basal ganglia. Nature 533, 402–406, doi:10.1038/nature17639 (2016).

Zhou, P. et al. Efficient and accurate extraction of *in vivo* calcium signals from microendoscopic video data. eLife, doi: 10.7554/eLife.28728 (2018).

